# Developmental dynamics of voltage-gated sodium channel isoform expression in the human and mouse neocortex

**DOI:** 10.1101/2020.11.18.389171

**Authors:** Lindsay Liang, Siavash Fazel Darbandi, Sirisha Pochareddy, Forrest O. Gulden, Michael C. Gilson, Brooke K. Sheppard, Atehsa Sahagun, Joon-Yong An, Donna M. Werling, John L.R. Rubenstein, Nenad Šestan, Kevin J. Bender, Stephan J. Sanders

**Author notes:** Please address correspondence to (S.J.S.).

## Abstract

**Objective:** Genetic variants in the voltage-gated sodium channels *SCN1A, SCN2A, SCN3A,* and *SCN8A* are leading causes of epilepsy, developmental delay, and autism spectrum disorder. The mRNA splicing patterns of all four genes vary across development in the rodent brain, including mutually exclusive copies of the fifth protein-coding exon detected in the neonate (5N) and adult (5A). A second pair of mutually exclusive exons is reported in *SCN8A* only (18N and 18A). We aimed to quantify the expression of individual exons in the developing human neocortex.

**Methods:** RNA-seq data from 176 human dorsolateral prefrontal cortex samples across development were analyzed to estimate exon-level expression. Developmental changes in exon utilization were validated by assessing intron splicing. Exon expression was also estimated in RNA-seq data from 58 developing mouse neocortical samples.

**Results:** In the mature human neocortex, exon 5A is consistently expressed at least 4-fold higher than exon 5N in all four genes. For *SCN2A, SCN3A,* and *SCN8A* a synchronized 5N/5A transition occurs between 24 post-conceptual weeks (2^nd^ trimester) and six years of age. In mice, the equivalent 5N/5A transition begins at or before embryonic day 15.5. In *SCN8A,* over 90% of transcripts in the mature human cortex include exon 18A. Early in fetal development, most transcripts include 18N or skip both 18N and 18A, with a transition to 18A inclusion occurring from 13 post-conceptual weeks to 6 months of age. No other protein-coding exons showed comparably dynamic developmental trajectories.

**Significance:** Splice isoforms, which alter the biophysical properties of the encoded channels, may account for some of the observed phenotypic differences across development and between specific variants. Manipulation of the proportion of splicing isoforms at appropriate stages of development may act as a therapeutic strategy for specific mutations or even epilepsy in general.

## 1. Introduction

Genetic variation in the genes *SCN1A, SCN2A, SCN3A,* and *SCN8A* are a major cause of epileptic encephalopathy (EE), autism spectrum disorder (ASD), and developmental delay.^1–3^ These four homologous genes encode voltage-gated sodium channels (Na_v_1.1, Na_v_1.2, Na_v_1.3, and Na_v_1.6 respectively) that are critical for a range of functions in the central nervous system,^4^ including axonal action potential initiation and propagation,^5,6^ dendritic excitability,^7,8^ macroscopic anatomical development,^9^ and activity-dependent myelination.^10^ The functional role, subcellular location, expression-level, and isoform selection of voltage-gated sodium channels vary across development and understanding this relationship is critical for understanding the etiology of the associated disorders and their therapeutic management.^7,11–19^ While some isoform-level differences have been assayed in rodents and mature human brains,^20–22^ the trajectories in the developing human cortex have not been described.^23^

Sodium channel genes are composed of multiple exons, which can be protein-coding (CDS for CoDing Sequence), untranslated regions (UTRs), or non-coding exons (NCEs). Differing combinations of these exons are called isoforms, which can change the amino acid sequence of the encoded proteins (proteoforms). The best-characterized isoform change across these four sodium channels are the two mutually exclusive copies of the fifth protein-coding exon.^17,24^ This exon encodes part of the first domain of the Na_v_ channel, including the end of transmembrane segment S3, most of transmembrane segment S4, and a short extracellular linker connecting these two segments. In humans, each copy of this fifth protein-coding exon is 92 nucleotides in length, encoding 30 amino acids, of which one to three amino acids vary between the two exon copies for each gene (Fig. 1B). ‘A’ isoforms (5A) include the ancestral and canonical copy, with an aspartic acid residue (Asp/D) encoded at position 7 of 30.^25^ ‘N’ isoforms (5N) use the alternative copy, with an asparagine (Asn/N) residue at position 7 of 30 in *SCN1A, SCN2A,* and *SCN8A* and a serine residue (Ser/S) in *SCN3A.* Despite this relatively small change in protein structure, differential inclusion of 5N or 5A can have marked effects on channel function. Indeed, these splice isoforms can alter channel electrophysiological characteristics,^26,27^ the functional impacts of variants associated with seizure,^23^ neuronal excitability,^28^ response to anti-epileptics,^21,22,26^ and seizure-susceptibility.^28^

**Figure 1.**
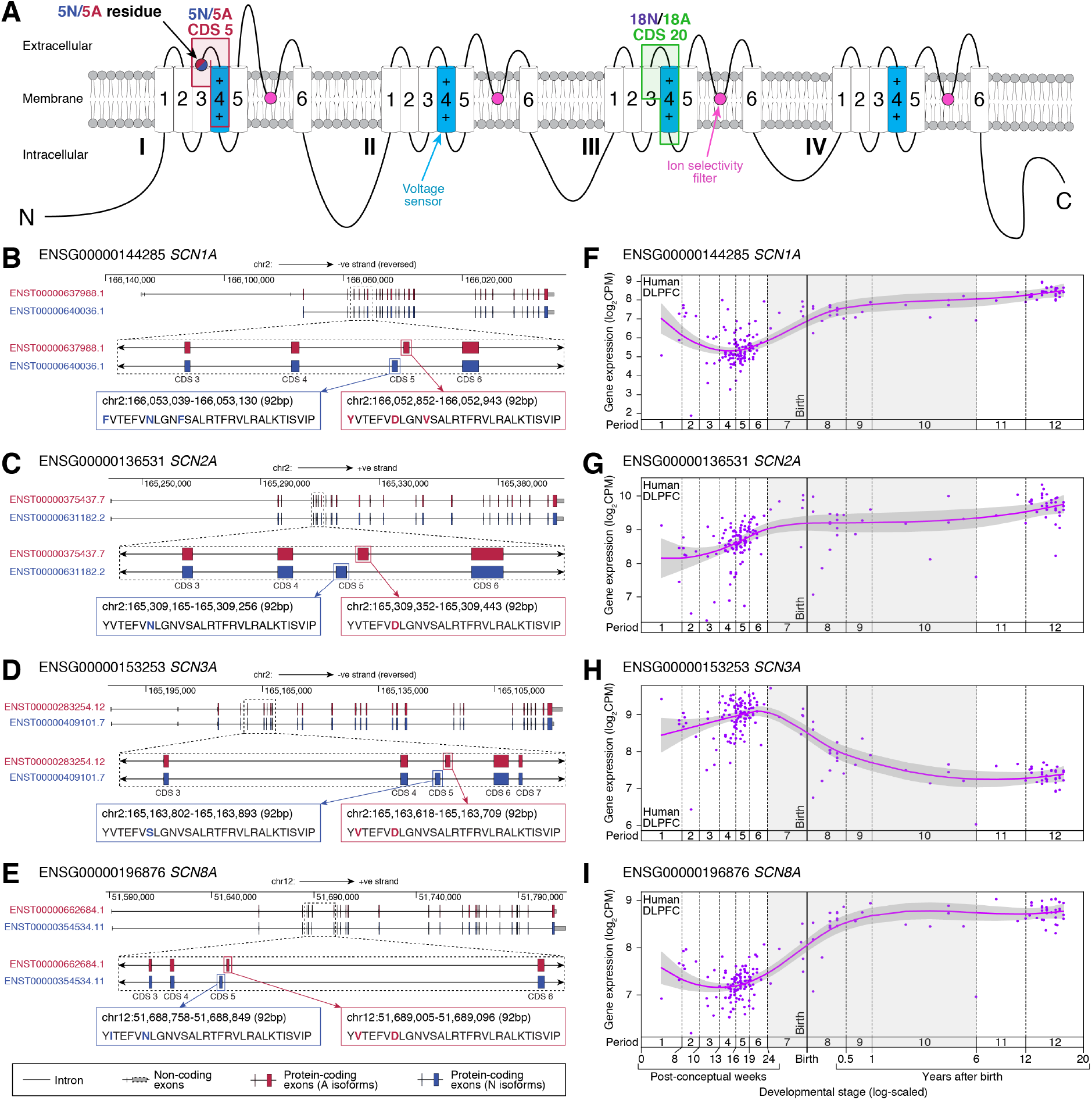
Splicing isoforms in voltage-gated sodium channels. **A)** Voltage-gated sodium channels are composed of four similar domains (I, II, III, IV), each of which includes six transmembrane segments with extracellular or intracellular linkers. The fourth transmembrane segment (S4) in each domain acts as a voltage sensor. Between the fifth and sixth transmembrane segment (S5, S6) is a pore loop that forms the ion selectivity filter. The fifth protein-coding exon (5N/5A, CDS 5) encodes a portion of the first domain, while the 20^th^ protein-coding exon (18N/18A, CDS 20) encodes a similar portion of the third domain. **B)** Location, genomic coordinates (GRCh38/hg38), and amino acid sequence of the ‘5A’ and ‘5N’ exons four sodium channels. **C)** Patterns of whole-gene expression in the human dorsolateral prefrontal cortex (DLPFC) across prenatal and postnatal development from the BrainVar dataset ^32^. CPM: counts per million. Genomic coordinates are based on GRCh38/hg38 using GENCODE v31 gene definitions.

The utilization of the 5N or 5A varies across development, with 5N generally being expressed at higher levels in the neonatal period while 5A predominates in adults.^27^ This switch is defined best in mouse, where the 5N:5A ratio varies by gene and brain region along with developmental stage.^20^ For *Scn2a* in mouse neocortex, the 5N:5A ratio is 2:1 at birth (postnatal day 0/P0) and flips to 1:3 by P15. For both *Scn3a* and *Scn8a,* 5A predominates throughout the postnatal period with a 1:2 ratio at P0 increasing to 1:5 by P15.^20^ *Scn1a* lacks a functional copy of 5N in the mouse genome. Similar developmental profiles currently have not been reported for humans beyond the of 5N/5A utilization *SCN1A* in adults, in which a 5N:5A ratio of over 1:5 was observed in the temporal cortex and hippocampus of adult surgical resections.^21,22^

In addition to the 5N/5A switch, a similar developmental shift in mutually exclusive exons has been reported for “exons 18N or 18A” in *SCN8A* only, regulated by the RNA-binding protein RBFOX1.^16,29,30^ Using GENCODE human v31 gene definitions,^31^ 18A maps to the 20^th^ proteincoding exon of major *SCN8A* isoforms (CDS 20, Fig. 1A), while 18N encodes the 8^th^ and last protein-coding exon (CDS 8) of a shorter eight protein-coding exon transcript (ENST00000548086.3, Fig. S1). In the embryonic mouse brain, most *SCN8A* transcripts include 18N or skip both 18N and 18A, leading to non-functional channels, while 18A predominates in the adult mouse and human brain.^16^

Here, we present data on the utilization of GENCODE-annotated protein-coding exons in four seizure-associated voltage-gated sodium channels in the human and mouse neocortex across development. We demonstrate a synchronized transition from 5N to 5A utilization between 24 post-conceptual weeks (2^nd^ trimester) and six years-of-age across all four voltage-gated sodium channels and a transition from 18N to 18A in *SCN8A* from 13 post-conceptual weeks to 6 months-of-age. These isoform differences can modify the function of the encoded voltagegated sodium channels, raising the potential that interventions, such as antisense oligonucleotides, could be used to modify the isoform ratio as a potential therapy for disorders caused by variants in sodium channel genes or epilepsy.

## 2. Materials and Methods

### 2.1 Genomic data

To quantify the relative proportion of protein-coding exon expression across development in the human cortex, we assessed bulk tissue RNA-seq data from 176 *post mortem* dorsolateral prefrontal cortex (DLPFC) samples from the BrainVar cohort.^32^ The BrainVar cohort also has corresponding whole-genome sequencing data that were used to derive per sample genotypes, as described previously.^32^ To assess corresponding patterns of exon expression in mouse cortex across development, we assessed 58 samples with bulk tissue RNA-seq data in wildtype C57/B6 mice. Thirty-four of these were generated as controls for ongoing experiments and 24 were downloaded from GEO.^33^

### 2.2 Exon expression

To assess exon expression in the human cortex, the 100bp paired-read RNA-seq data from BrainVar were aligned to the GRCh38.p12 human genome using STAR aligner^34^ and exon-level read counts for GENCODE v31 human gene definitions were calculated with DEXSeq^35^ and normalized to counts per million (CPM).^36^ Despite the similar amino acid sequence, the nucleotide sequence of 5N and 5A is sufficiently differentiated across the four genes that 100bp reads align unambiguously to one location in the genome.^37^ Reads were detected in 5N and 5A for all samples, across all four genes, with the exception of the *SCN1A* for which 31 of 176 samples (17.6%) had no detectable 5N reads (Fig. 2A). Along with quantifying the expression of 5N and 5A (Fig. 2), we also assessed expression for the surrounding constitutive exons, as a control (Fig. S2). For the mouse cortical data, the same analysis methods were used but with alternative references, specifically the GRCm38/mm10 genome and GENCODE vM25 gene definitions. A similar approach was used to assess the utilization of 18N and 18A in *SCN8A.*

**Figure 2.**
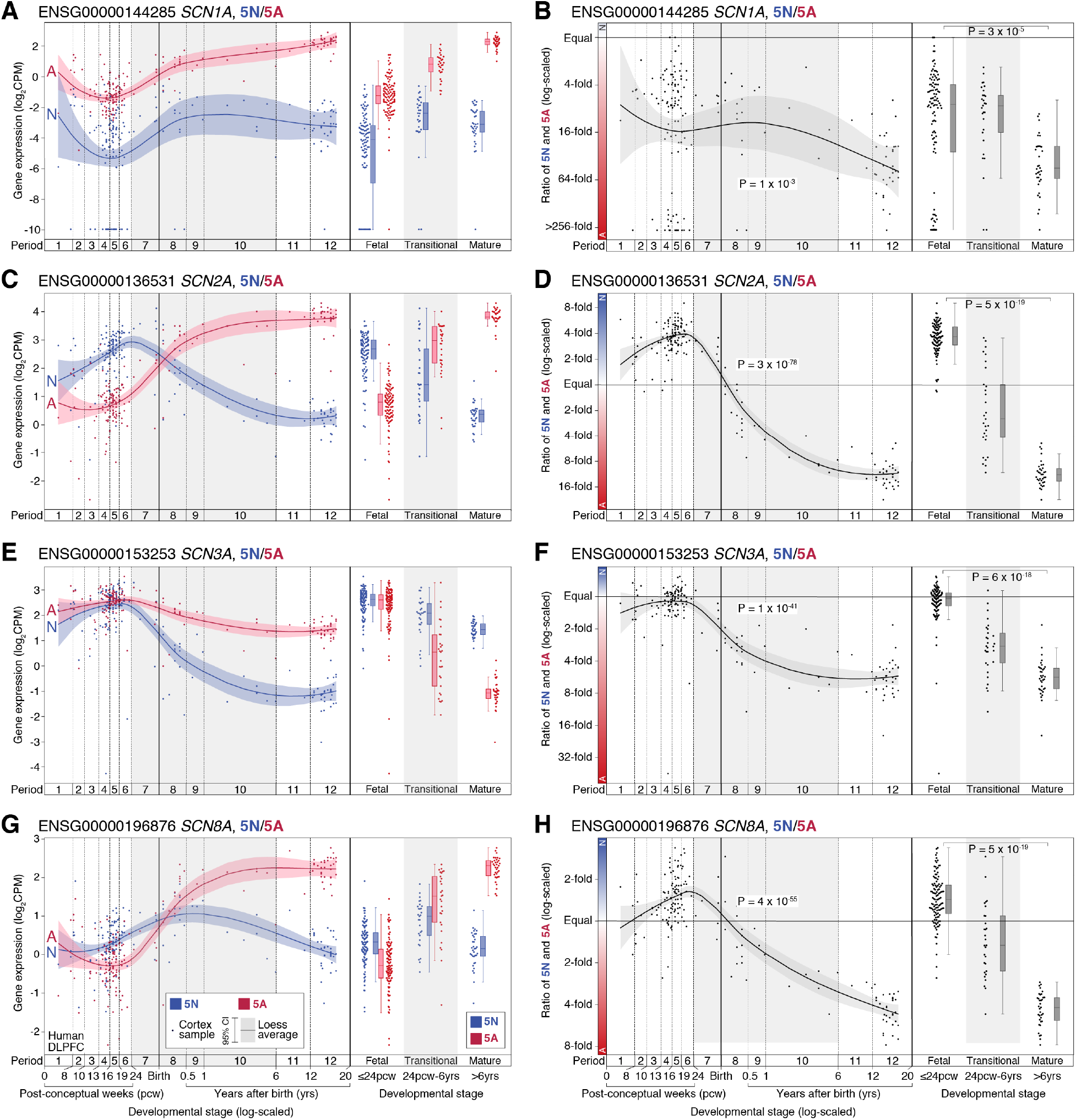
Expression of 5N and 5A in the human cortex across development. **A)** The expression of 5N (blue) and 5A (red) in *SCN1A* is shown for 176 BrainVar human cortex (DLPFC) samples across development (points). On the left, the colored line shows the Loess smoothed average and 95% confidence interval (shaded region). On the right, boxplots show the median and interquartile range for the same data, binned into fetal, transitional, and mature developmental stages. **B)** The ratio of 5N and 5A expression from panel ‘A’ is shown across development (left) and in three developmental stages (right). **C-H)** Panels A and B are repeated for the genes *SCN2A, SCN3A, SCN8A.* For comparison, the same plots for CDS four and six are shown in Figure S2. CPM: Counts per million; DLPFC: Dorsolateral prefrontal cortex. Statistical tests: B, D, F, H) Left panel, linear regression of log_2_(5N:5A ratio) and log_2_(post-conceptual days). Right panel, two-tailed Wilcoxon test of log_2_(5N:5A ratio) values between fetal and mature groups.

### 2.3 Intron splicing

We applied a complementary approach to detecting 5N and 5A exon usage by assessing intron splicing via reads that map across exon-exon junctions in the same 176 BrainVar samples. Reads were aligned with OLego aligner^38^ using the same genome build and gene definitions as for exon expression. Clusters of differential intron splicing were identified with Leafcutter^39^ and differences across development were detected by comparing 112 prenatal samples to 60 postnatal samples. No cluster was detected for 5N/5A in *SCN1A,* preventing assessment across development, but clusters were identified and assessed for the other three genes and for 18N/18A in *SCN8A* (Figs. 3, 5).

**Figure 3.**
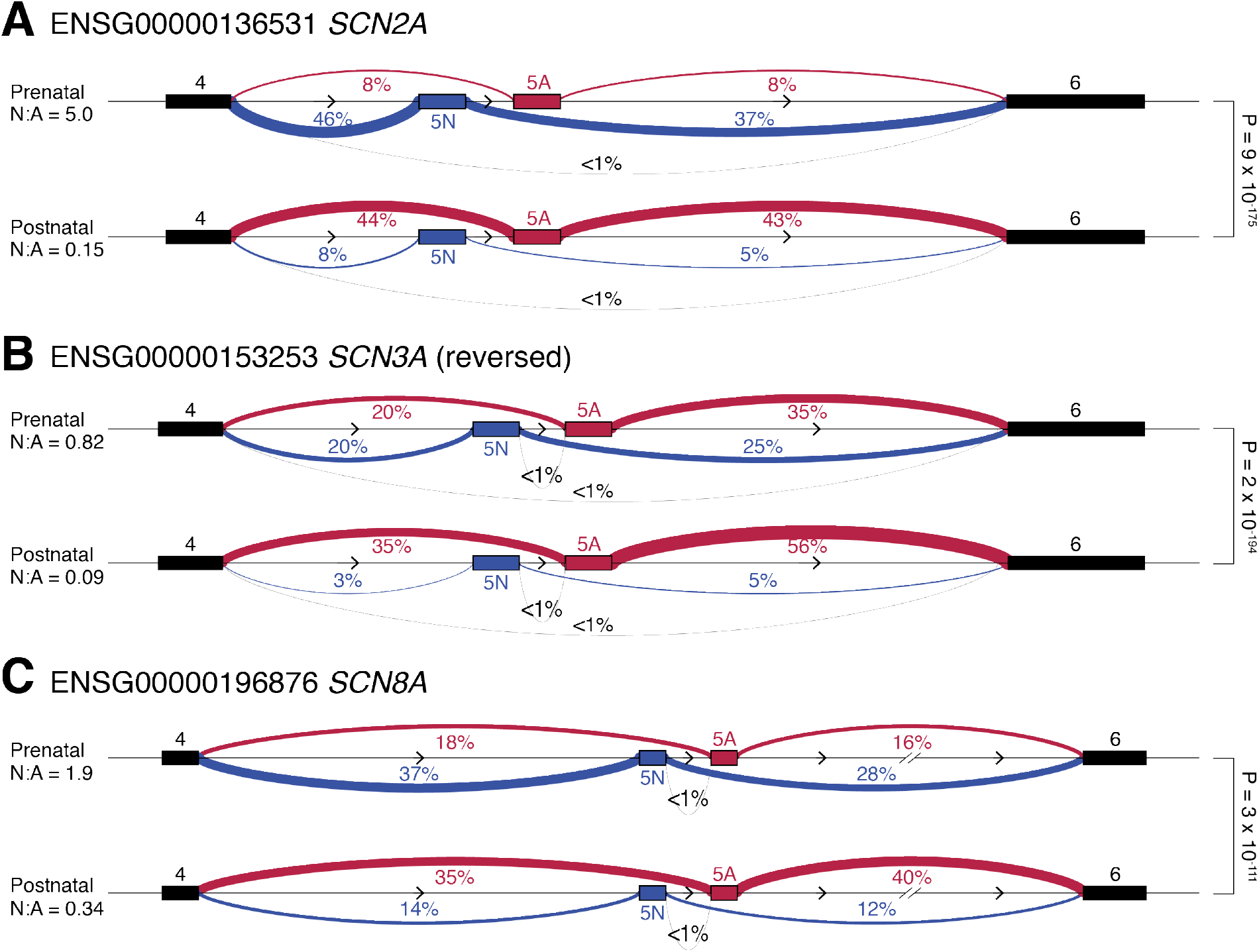
Intron splicing of sodium channel genes in the developing human cortex. **A)** Sashimi plot of splicing in the prenatal (top, N=112 samples) and postnatal (bottom, N=60 samples) DLPFC for *SCN2A.* Linewidth is proportional to percentage of split reads observed for each intron and this value is given as a percentage. Introns related to 5N inclusion are shown in blue, those related to 5A inclusion are shown in red, and others are in grey. **B-C)** Equivalent plots for *SCN3A* (a negative strand gene with the orientation reversed to facilitate comparison to the other two genes) and *SCN8A.* P-values compare the prenatal and postnatal cluster using a Dirichlet-multinomial generalized linear model, as implemented in Leafcutter.^39^

### 2.4 Quantitative trait locus (QTL) analysis

Common variants with a minor allele frequency ≥5% in both the prenatal (N = 112) and postnatal (N = 60) samples and Hardy Weinberg equilibrium p value ≥ 1×10^-12^ were identified previously.^32^ Variants within one million basepairs of each sodium channel gene were extracted and integrated with the Leafcutter clusters, along with the first five principal components calculated from common variants identified in whole-genome sequencing data from these samples and 3,804 parents from the Simons Simplex Collection^32,40^ to predict sQTLs with FastQTL.^41^ This analysis was performed on all samples, prenatal-only samples, and postnatal-only samples, with false discovery rate (FDR) estimated from the results of each analysis using the Benjamini-Hochberg procedure.^42^ To assess correlation of 5N expression for the SNP rs3812718, genotypes were extracted for chr2:166,053,034 C>T (GRCh38) and compared with 5N expression calculated by DEXSeq, as described above.

### 2.5 Statistical analysis

The 5N:5A expression ratio was calculated from normalized exon expression values (CPM). Linear regression was used to assess whether this ratio varied across development by comparing the log-transformed 5N:5A ratio to log-transformed post-conceptual days (Fig. 2). The difference in ratio was also assessed between the mid-late fetal samples (N=112) and childhood/adolescent/young adult samples (N=35) with a two-tailed Wilcoxon test. To compare intron splicing between prenatal and postnatal samples, we used the P-values estimated with a Dirichlet-multinomial generalized linear model, as implemented in Leafcutter.^39^

## 3. Results

### 3.1 Expression of voltage-gated sodium channels in the human cortex

Gene expression varies dramatically across development for many genes, especially during the late-fetal transition, during which half the genes expressed in the brains undergo a concerted increase or decrease in expression.^12,32,43,44^ To assess gene-level developmental trajectories, we analyzed bulk-tissue RNA-seq of the human DLPFC in 176 *post mortem* samples from the BrainVar cohort (104 male, 72 female, spanning 6 post-conceptual weeks to 20 years after birth).^32^ The gene-level expression profile of all four voltage-gated sodium channels changes during this late-fetal transition (Fig. 1F-I), with *SCN1A, SCN2A,* and *SCN8A* expression rising from mid-fetal development through infancy to early childhood, while *SCN3A* expression falls.

### 3.2 Developmental trajectories of 5N and 5A expression in the human cortex

The majority of protein-coding exons follow the expression trajectory of their parent gene across development (Fig. S3), however all four sodium channels show dynamic changes in the utilization of 5N/5A (Fig. 2). This is especially marked for *SCN2A* and *SCN8A,* where 5N is expressed at a higher level than 5A in the mid-fetal brain but this reverses soon after birth. Plotting the 5N:5A ratio allows exon utilization to be assessed independent of changes in gene expression (Fig. 2). All four genes show changes in the N:A ratio across development, with a modest change for *SCN1A* (0.14 fetal to 0.02 childhood/adolescent; p=0.00003, two-sided Wilcoxon test, Figure 2B) and dramatic changes for *SCN2A* (3.7 to 0.09; p=5×10^-19^, Fig. 2D), *SCN3A* (0.96 to 0.18; p=6×10^-18^, Fig. 2F), and *SCN8A* (3.7 to 0.09; p=5×10^-19^, Fig. 2H). As a control, we applied this approach to assess the ratio of CDS 4 and CDS 6 across development. We observed no developmental shift in the 4:6 ratio for *SCN1A, SCN2A,* and *SCN3A,* however the exon 4:6 ratio is marginally higher than expected in the prenatal period for *SCN8A* (0.82 vs. 0.66; 9×10^-10^, Fig. S2). This developmental variation in *SCN8A* is not observed for the surrounding protein-coding exons and reflects a modest increase in CDS 4 expression in the prenatal period, based on the expected expression given the exon length (Fig. S4).

### 3.3 Intron splicing around 5N and 5A in the human cortex

To verify that mutually exclusive use of 5N and 5A underlies the observed exon expression changes (Fig. 2), we considered RNA-seq reads that spanned exon-exon junctions to quantify intron splicing. Clusters of differential intron splicing corresponding to 5N/5A usage were identified by Leafcutter for *SCN2A, SCN3A,* and *SCN8A* (Fig. 3), but not *SCN1A,* likely due to the consistently low expression of N isoforms (Fig. 2). The splicing patterns for *SCN2A, SCN3A,* and *SCN8A* are consistent with the observed exon expression changes (Fig. 2, 3) and at least 99% of reads are consistent with mutually exclusive 5N/A utilization.

### 3.4 Developmental trajectories of 5N and 5A expression in the mouse cortex

We repeated the analysis of sodium channel 5N/5A expression using bulk tissue RNA-seq data from the mouse cortex across development (N=58; E15.5 to P75). Our data are consistent with the N:A ratios described previously.^20^ We observe more substantial differences at the extremes of development: *SCN2A* (3.3 fetal to 0.06 mature; p=0.00003, two-sided Wilcoxon test, Fig. 4C), *SCN3A* (2.4 to 0.14; p=0.00008, Fig. 4E), and *SCN8A* (1.8 to 0.22; p=0.00003, Fig. 4G). Mice lack a functional 5N exon in *SCN1A.*

**Figure 4.**
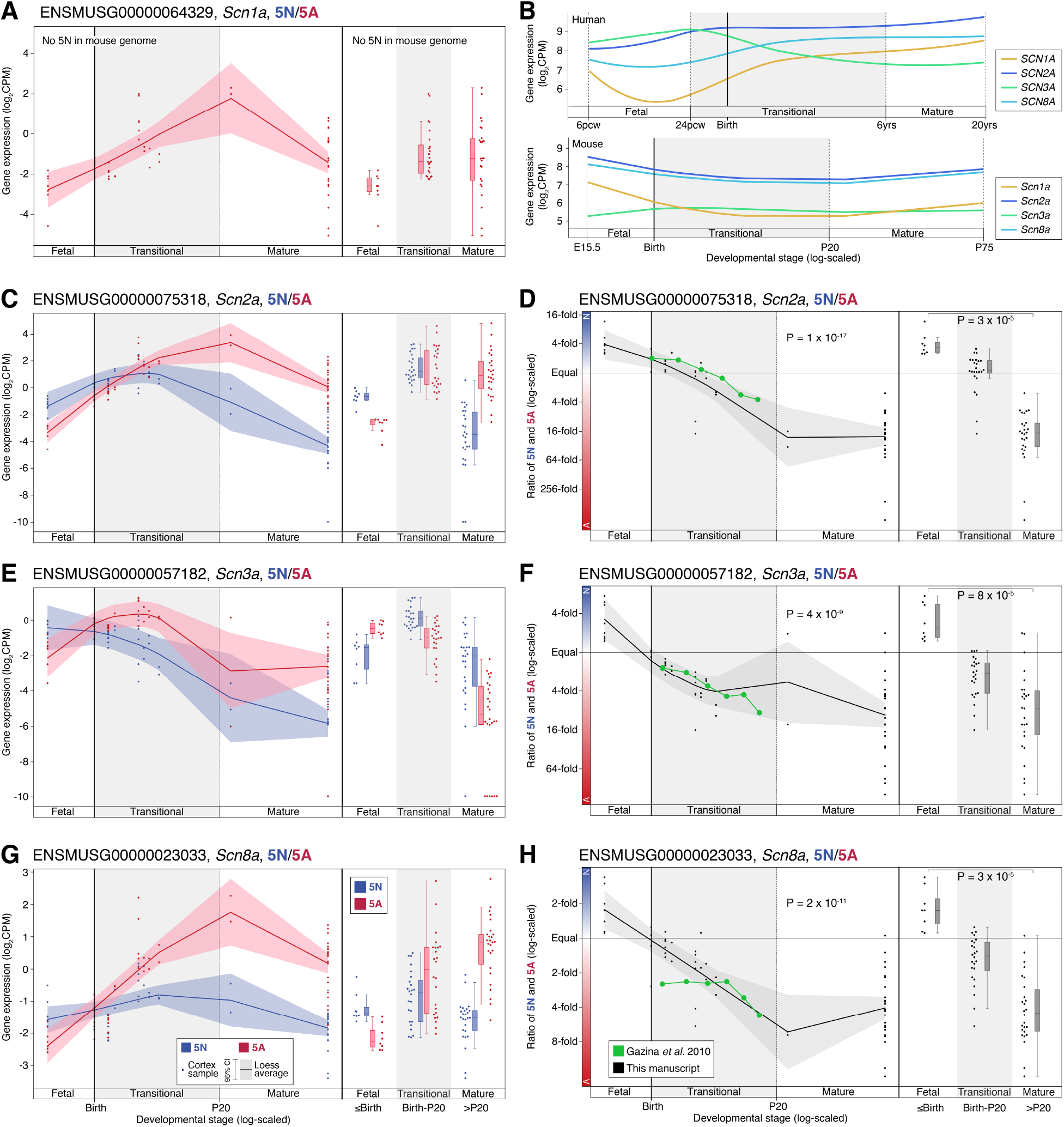
Expression of 5N and 5A in the mouse cortex across development. **A)** The expression of 5A (red) in *Scn1a* is shown for 58 mouse cortex samples across development (points); no functional 5N equivalent is present in the mouse genome. On the left, the colored line shows the Loess smoothed average and 95% confidence interval (shaded region). On the right, boxplots show the median and interquartile range for the same data, binned into fetal, transitional, and mature developmental stages. **B)** The Loess smoothed average expression of the four voltage-gated sodium channels in human cortex (top, Fig. 1) and mouse cortex (bottom). **C)** Panel ‘A’ is repeated for *Scn2a* with the inclusion of 5N (blue). **D)** The ratio of 5N and 5A expression from panel ‘C’ is shown across development (left) and in three developmental stages (right). Values reported previously in mouse cortex are shown in the same scale in green for comparison ^20^. **E-H)** Panels ‘C’ and ‘D’ are repeated for the genes *Scn3a, Scn8a.* CPM: Counts per million. Statistical tests: D, F, H) Left panel, linear regression of log_2_(5N:5A ratio) and log_2_(post-conceptual days). Right panel, two-tailed Wilcoxon test of log_2_(5N:5A ratio) values between fetal and mature groups.

### 3.5 No evidence of common polymorphisms regulating 5N or 5A utilization

A common polymorphism (rs3812718, GRCh38 chr2:166,053,034 C>T, IVS5N+5G>A) has previously been associated with epilepsy, seizures, and response to anti-epileptics,^21,22,26,45,46^ though this variant did not reach genome-wide significance in a mega-analysis of epilepsy.^47^ Prior analyses of expression in the adult human temporal cortex showed evidence that the homozygous variant allele (TT in DNA, AA in cDNA) was associated with reduced utilization of 5N.^21,48^ We do not observe evidence for such a relationship in the prenatal or postnatal prefrontal cortex (Fig. S5) and this polymorphism is not identified as a splicing quantitative trait locus (QTL) in GTEx.^49^ Furthermore, this variant is not predicted to alter splicing behavior using the SpliceAI algorithm.^50^ The TT genotype is associated with increased expression of *SCN1A* in the adult human basal ganglia with (p=1×10^-10^).^49^

### 3.6 Developmental trajectories of 18N and 18A expression in *SCN8A*

We next considered the developmental timing of the transition between 18N and 18A in *SCN8A* (Fig. 1A, 5A). Intron splicing shows a robust difference between prenatal and postnatal human dorsolateral prefrontal cortex (P = 4 x 10^-185^, Fig. 5B), with the prenatal period characterized by high frequencies of transcripts excluding 18A, either including 18N or skipping both 18N and 18A, while in the postnatal cortex 18A is included in 93% of reads. Considering exon expression (Fig. 5C, 5D), the expression of 18A increases markedly over development and this is distinct from other protein-coding exons for *SCN8A* (Fig. S3). The 18N/18A transition begins around 13 post-conceptual weeks and continues till six months-of-age, with both timepoints being earlier than the equivalents for 5N/5A in *SCN8A* and the other genes.

**Figure 5.**
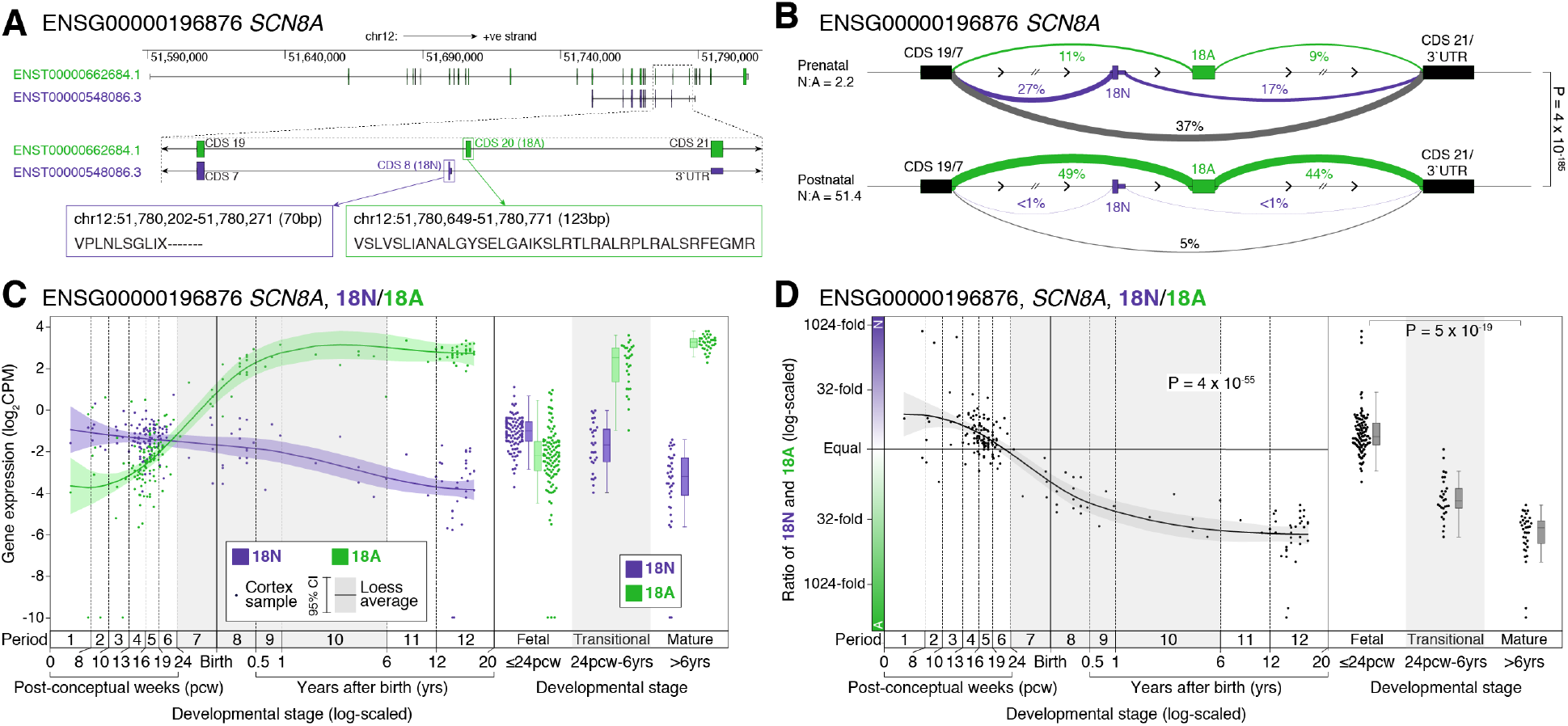
Developmental trajectories of CDS 20 (18N/18A) in human cortex in *SCN8A.* **A)** Location, genomic coordinates (GRCh38/hg38), and amino acid sequence of the 18N and 18A exons in *SCN8A.* **B)** Sashimi plot of intron splicing in the prenatal (top, N=112 samples) and postnatal (bottom, N=60 samples) dorsolateral prefrontal cortex. Linewidth is proportional to percentage of reads observed for each exon-exon junction and this value is also shown as a percentage. Introns related to 18N exon inclusion are shown in purple, those related to 18A exon inclusion are shown in green, and others are in grey. **C)** Expression of the 18N (purple) and 18A (green) for 176 BrainVar human dorsolateral prefrontal cortex samples across development (points). On the left, the colored line shows the Loess smoothed average with the shaded area showing the 95% confidence interval. On the right, boxplots show the median and interquartile range for the same data, binned into fetal, transitional, and mature developmental stages. **D)** The 18N:18A ratio is shown for each sample from panel ‘C’ across development (left) and binned into three groups (right). CPM: Counts per million; Statistical analyses: B) Dirichlet-multinomial generalized linear model, as implemented in Leafcutter,^39^ D) Left panel, linear regression of log_2_(18N:18A ratio) and log_2_(post-conceptual days). Right panel, two-tailed Wilcoxon test of log_2_(18N:18A ratio) values between fetal and mature groups.

### 3.7 Other annotated protein-coding exons with distinct developmental trajectories

To assess whether other protein-coding exons undergo distinct developmental transitions (Fig. S3), we calculated the ratios of all pairs of protein-coding exons within each for the four sodium channel genes and assessed whether the ratio was correlated with development stage using linear regression. This is the same calculation used to quantify the 5N/5A and 18N/18A transitions (Fig. 2, 5D) and distinguishes exons with expression profiles that differ from the rest of the gene (e.g. 5A in *SCN2A),* rather than simply being expressed at reduced levels, suggesting alternative regulatory processes (Fig. S3). Visualizing the R^2^ values of these correlations provides simple method to identify the such distinct trajectories (Fig. 6). Aside from 5N/5A and, in *SCN8A,* 18N/18A, no protein-coding exons common to most isoforms (consistent CDS in Fig. S3) show differential expression, but a few weakly expressed protein-coding exons specific to a small number of isoforms (variable CDS in Fig. S3) do vary across development (Fig. 6).

**Figure 6.**
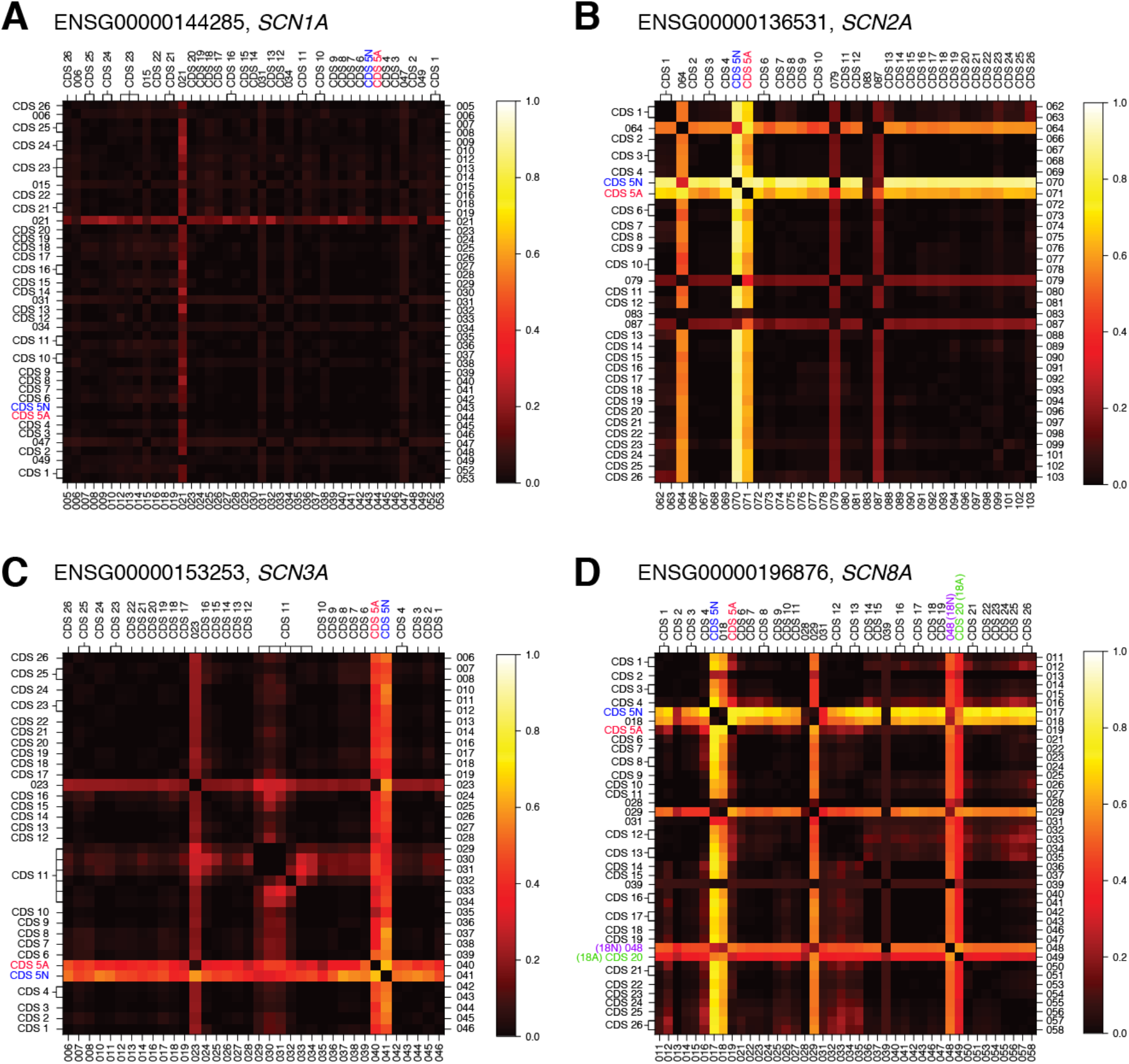
Identification of protein-coding exons with complex developmental trajectories. **A)** The correlation between the ratio of CPM expression between pairs of exons (log-scaled) and developmental stage (post conceptual days, log-scaled) for *SCN1A* was assessed with a linear model (e.g. Fig. 2B). The R^2^ value of each exon pair is show as a heat map with ‘hot’ colors representing exon-pairs with high R^2^ values for which variation in the ratio is correlated with developmental age, i.e. pairs of exons that show substantially different expression across development. Exon numbers from DEXSeq (Table S2) are shown on the bottom and right and equivalent CDS number on the top and left (see Table S2). **B-D)** The analysis is repeated for *SCN2A, SCN3A,* and *SCN8A.*

GENCODE defines seven variable CDS exons for *SCN1A* (DEXSeq divisions: 006, 015, 021, 031, 034, 047, 049; Table S2, Fig. 6A). Of these, only 021 shows a distinct developmental trajectory (Fig. 6A), with reduced postnatal expression relative to other *SCN1A* exons (Fig. S3). This result is verified by the intron splicing data (p = 6 x 10^-91^, Leafcutter).

In *SCN2A,* the 5N/5A trajectories stand out clearly (Fig. 6B). There are four variable CDS exons (DEXSeq divisions: 064, 079, 083, 087; Table S2, Fig. 6B), three of which have distinct developmental trajectories (Fig. 6B, S3): 064 (Fig. S3, P = 2 x 10^-12^, Leafcutter), 079 (Fig. S3, P = 7 x 10^-33^, Leafcutter), 087 (Fig. S3, P = 2 x 10^-20^, Leafcutter). The single variable CDS exon in *SCN3A,* 023 (Table S2, Fig. 6C), varies across development (Fig. S3, P = 3 x 10^-80^, Leafcutter).

Finally, aside from 18N, there are five variable CDS exons in *SCN8A* (DEXSeq divisions: 018, 028, 029, 031, 039; Table S2, Fig. 6D) of which 018 and 029 vary across development (Fig. 6D), but neither of these are validated by Leafcutter.

## Discussion

Using transcriptomic data from 176 human dorsolateral prefrontal cortex samples, we characterized the developmental patterns for all protein-coding exons in *SCN1A, SCN2A, SCN3A,* and *SCN8A* (Fig. 6, S3). We observed a coordinated decrease in the 5N:5A ratio between 24 post-conceptual weeks (2^nd^ trimester) and six years-of-age that is synchronized with widespread transcriptomic changes in the brain during the late-fetal transition.^32,44^ This is preceded by a similar decrease in the 18N:18A ratio in *SCN8A* from 13 post-conceptual weeks to 6 months-of-age, which is regulated by *RBFOX1.* By analyzing a wider developmental window than prior analyses^20,21,48^ we observed more dynamic changes and larger disparities in exon expression.

Recent advances have shown that differential splicing patterns can be effective therapeutic targets in humans, for example through intrathecal antisense oligonucleotides.^51,52^ Since the electrophysiological consequences of some epileptic encephalopathy associated variants differ between 5N and 5A, manipulating this ratio may offer therapeutic benefit in individuals carrying these variants. We consider three therapeutic scenarios.

First, for individuals with disorder-associated genetic variants within the 30 amino acids encoded by the 5^th^ exon, expressing the other copy of the 5^th^ exon could skip the variant. Theoretically, this approach could benefit individuals with both loss-of-function (protein-truncating variants, missense, splice site) and gain-of-function (missense) variants in the 5^th^ exon. At present, ten such cases have been identified, all with epileptic encephalopathy variants identified in the 5A exon of *SCN2A* and *SCN8A*.^53,54^ Since the total transcript level would be unchanged, this strategy may provide a wider therapeutic window than simply decreasing expression levels. The success of the therapy would depend upon the proportion of transcripts expressing the alternate 5^th^ exon and the ability of this exon to functionally replace the original 5^th^ exon.

Second, splice isoforms can also have an effect on the biophysical effects of variants outside the 5^th^ exon. For example, three recently characterized epileptic encephalopathy associated variants in *SCN2A*—T236S, E999K, and S1336Y—all exhibit more pronounced alterations in their electrophysiological properties in 5N NaV1.2 isoforms.^23^ Two other epileptic encephalopathy-associated variants—M252V and L1563V—exhibit biophysical changes only when expressed on 5N isoform.^14,55^ For individuals with these mutations, tilting expression towards the 5A isoform could provide some symptomatic improvement, especially during infancy. Including both the 5N and 5A isoforms in functional characterization of variant impact may identify many more such variants.^23^

Finally, modifying splicing might aid seizure control in older children and adults. At this age, the 5A isoform is predominantly utilized in both *SCN2A* or *SCN8A,* which are mainly expressed in glutamatergic neurons.^11^ Reverting expression to the fetal/neonatal state by encouraging 5N utilization could reduce the excitability of cortical glutamatergic neurons, potentially limiting seizures. Since this would require repeated intrathecal administration, it would likely be limited to the most severe cases of epilepsy. Furthermore, it remains to be seen whether this approach could offer therapeutic benefits above and beyond existing antiepileptic drugs.

Our analysis was limited by the use of short-read transcriptomic data, leading us to focus on quantifying exon-level expression (Fig. 2) and splice junction usage (Fig. 3), rather than relying on estimates of isoform utilization (Fig. S1). We also elected to focus on protein-coding transcripts and exons defined by GENCODE (v31) rather than attempting *de novo* transcriptome assembly. Emerging long-read transcriptomic technology may substantially expand estimates of isoform and exon diversity but these technologies have not been applied to the developing human brain at scale.^56,57^ We also note that transcriptomic data is only partially predictive of protein levels and other factors, including channel transport and degradation, may influence the impact of isoforms on neuronal function. Comparing the human and mouse cortex data (Figs. 2, 4), it is possible that more substantial differences in gene and exon expression may be observed at earlier embryonic times in the mouse or with larger sample sizes. In addition, the use of bulk-tissue transcriptomic data limits our ability to assess how individual cell-types or cell-states contribute to the observed isoform trajectories. Technological and methodological advances may provide insights at cell-level resolution in the future.^58^

## Conclusion

Dramatic differences in exon usage of *SCN1A, SCN2A, SCN3A,* and *SCN8A* observed in rodent brains also occur in the human developing cortex, beginning in mid-fetal development and continuing through childhood. These changes in splicing affect the biophysical properties of the encoded channels and are likely to contribute to differences in phenotype observed between individuals with different variants and across development.

## Supporting information

Table_S1_Samples.xlsx

Table_S2_Exons.xlsx

## Acknowledgements

This work was supported by funding provided by the Simons Foundation Autism Research Initiative (SFARI) grants 574598 (to S.J.S.), 647371 (to S.J.S.), 629287 (to K.J.B.), and 513133 (to K.J.B.), the National Institute for Mental Health (NIMH) grants: R01 MH111662 (to S.J.S.), U01 MH122681 (to S.J.S.), P50 MH106934 (to N.S.), R01 MH109904 (to N.S.), R01 MH110926 (to N.S.), and U01 MH116488 (to N.S.), the National Institute of Neurological Disorders and Stroke (NINDS) grant: R01 NS099099 (to J.L.R.R.), and the National Research Foundation of Korea: NRF-2020R1C1C1003426 (to J.Y.A.) and NRF-2017M3C7A1026959 (to J.Y.A.).

## Author Contributions

Experimental design, S.J.S.; Data generation, L.L., S.F.D., S.P., F.O.G., A.S., J.Y.A., and J.L.R.R.; Data processing, L.L., M.C.G., B.K.S., and D.M.W.; Data analysis, L.L., D.M.W., and S.J.S.; Statistical analysis, S.J.S.; Manuscript preparation, L.L., K.J.B., and S.J.S.

