## Supplemental_Text for "Developmental dynamics of voltage-gated sodium channel isoform expression in the human and mouse neocortex"

### Supplementary Materials

#### Data Acquisition

STAR aligned BAMs from the 176 Brainvar bulk RNA-seq samples were obtained from the PsychENCODE Knowledge Portal: syn21557948 on Synapse.org (https://www.synapse.org/#!Synapse:syn4921369), as described previously.[32] For the mouse data, FASTQ files for the mouse data for development periods before P28 were generated as controls for ongoing experiments. FASTQ files for samples at after P28 were obtained from NCBI's Sequence Read Archive[33] (Table S1). Sample ages for the BrainVar samples, expressed as post-conceptual days were obtained from the supplementary materials.[32]

#### RNA-seq alignment and exon-level read count quantification

For the mouse data, RNA-seq short reads were aligned with the STAR aligner to the Mus musculus reference GRCm38.p6 to generate BAM files matching the BrainVar data. For both the BrainVar and mouse datasets, exon level read counts for the genes *SCN1A*/*Scn1a* (ENSG00000144285.20/ENSMUSG00000064329.13), *SCN2A*/*Scn2a* (ENSG00000136531.16/ENSMUSG00000075318.13), *SCN3A*/*Scn3a* (ENSG00000153253.18/ENSMUSG00000057182.15), and *SCN8A*/*Scn8a* (ENSG00000196876.16/ENSMUSG00000023033.14) were calculated using DEXSeq,[35] annotating to GENCODE v31 of the human reference genome GRCh38 in humans and GENCODE vM25 of the mouse reference genome GRCm38 in mice. GENCODE GTF files were converted to GFF files using the DEXSeq package script 'dexseq_prepare_annotation.py'. Counts were acquired from the 'dexseq_count.py script'. Human samples were processed as paired-end and reverse stranded (-p yes -s reverse), while mouse samples were processed with a combination of paired and single endedness, and stranded and unstrandedness (Table S1). Raw read counts were converted into counts per million (CPM).[36]

#### Transcript level quantification

BAMs generated for the BrainVar dataset were converted to paired end FASTQ files using bedtools v.2.29.2 bamtofastq command.[59] Transcripts were quantified with 'salmon quant -i transcript_idx -l A -1 sample_r1.fastq.gz -2 sample_r2.fastq.gz -o output --validateMappings --gcBias --dumpEq' with version 1.3.0 of the software.[60] The transcript index was generated using all GRCh38 transcript sequences from GENCODE v31.

#### Exon junction quantification

Short reads from the BrainVar dataset were aligned to the human reference genome GRCh38 using OLego v.1.1.5.[38] The resulting BAMs were converted into junction files and run through Leafcutter's suite of tools for quantifying RNA-splicing variation.[39] Introns found in the junction files were clustered together with the default setting of 50 reads per cluster and a maximum intron length of 500kb, creating a matrix describing the read count per sample at a particular intron cluster site using the default parameters to the 'leafctter_cluster.py' script. The file used to define exons was derived from the GENCODE v.31 GTF for GRCh38 and is hosted in the Leafcutter git repo. Then, differential splicing analysis was run based on whether samples were identified as prenatal or postnatal using the default parameters to the 'leafcutter_ds.R' script.

The effect sizes outputted from the differential splicing analysis was processed with Leafcutter's 'Leafviz' visualization tool to annotate introns.

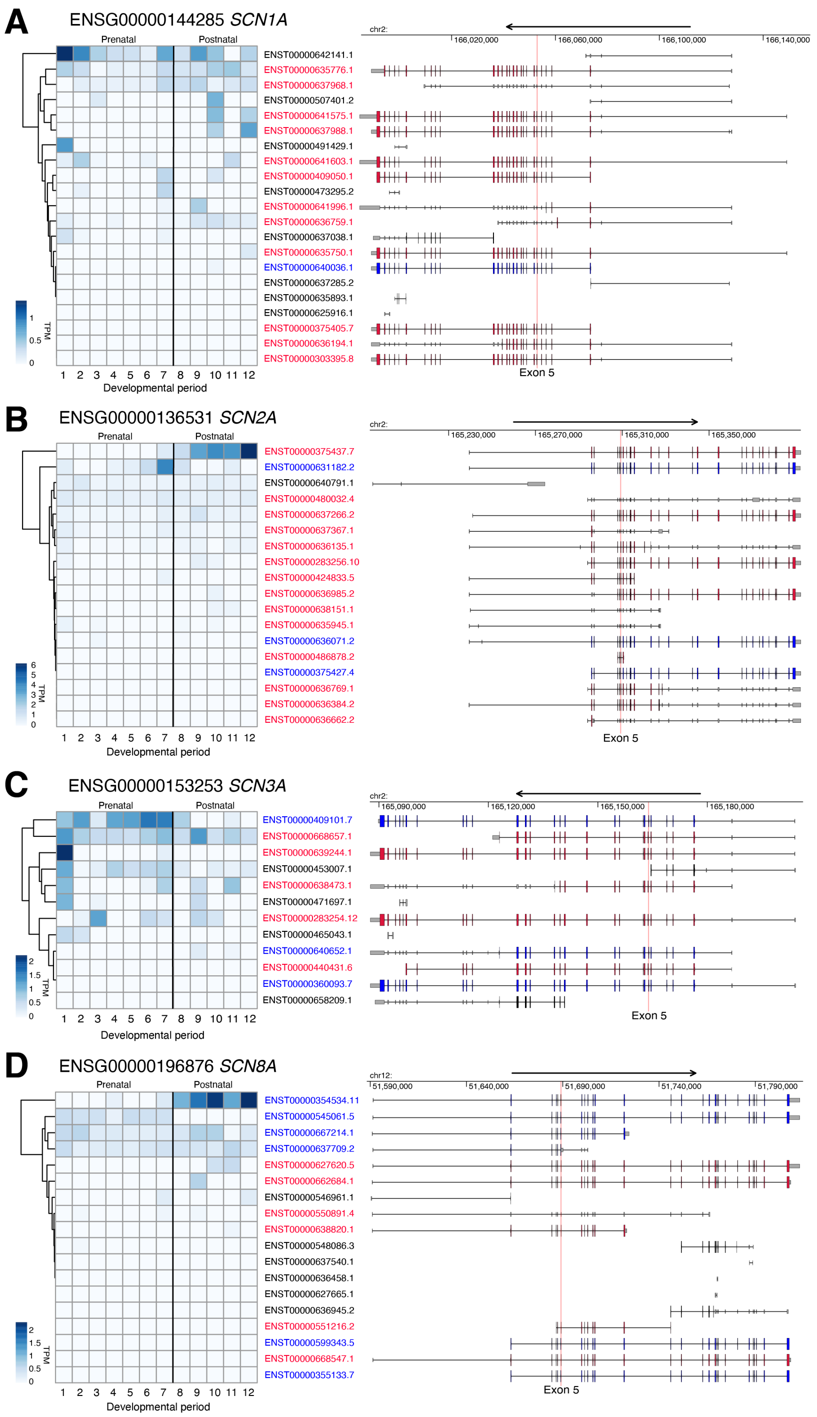


**Figure S1. Isoform usage in voltage-gated sodium channels predicted by Salmon. A)** For *SCN1A*, the estimated transcripts per million (TPM) values for each transcript in the human dorsolateral prefrontal cortex (DLPFC) across twelve developmental stages are represented by the shade of blue in the heatmap, with transcripts with high estimated expression in darker shades. N isoform transcripts that use exon 5N are shown in blue, A isoform transcripts (exon 5A) in red, and transcripts that do not include exon 5 are shown in black. Transcript definitions are from GENCODE v31 and genomic coordinates are shown in GRCh38. **B-D)** The representation in 'A' is repeated for *SCN2A*, *SCN3A*, and *SCN8A*.

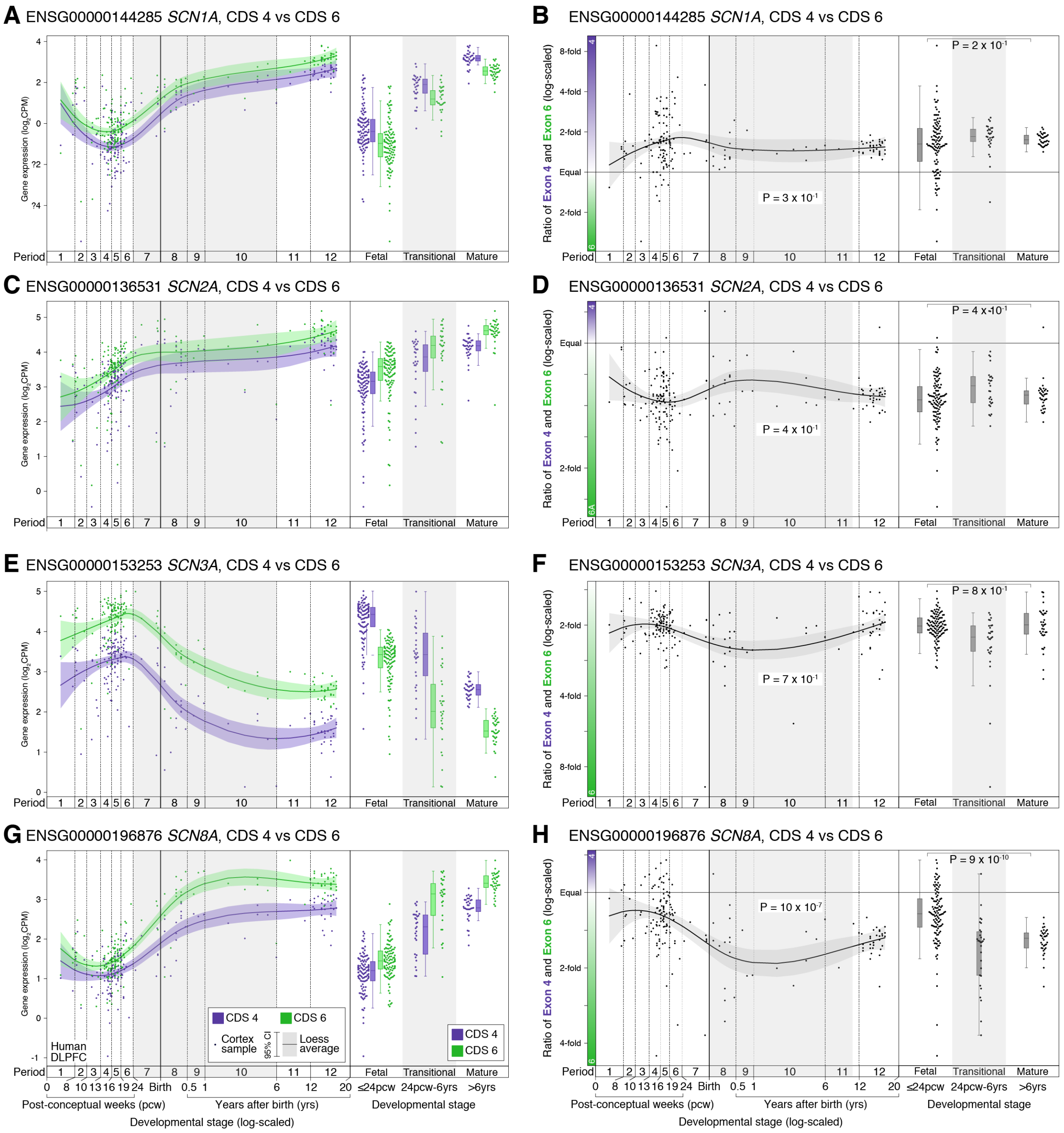


**Figure S2. Expression of CDS 4 and CDS 6 in the human cortex across development. A)** The expression of CDS 4 (purple) and CDS 6 (green) in *SCN1A* is shown for 176 BrainVar human dorsolateral prefrontal cortex samples across development (points). On the left, the colored line shows the Loess smoothed average and 95% confidence interval (shaded region). On the right, boxplots show the median and interquartile range for the same data, binned into fetal, transitional, and mature developmental stages. **B)** The ratio of CDS 4 and CDS 6 expression from panel 'A' is shown across development (left) and in three developmental stages (right). **C-H)** Panels A and B are repeated for the genes *SCN2A*, *SCN3A*, *SCN8A*. For comparison, the same plots for CDS four and six are shown in Figure S4. CPM: Counts per million. Statistical tests: B, D, F, H) Left panel, linear regression of $\log_2$(4:6 ratio) and $\log_2$(post-conceptual days). Right panel, two-tailed Wilcoxon test of $\log_2$(4:6 ratio) values between fetal and mature groups.

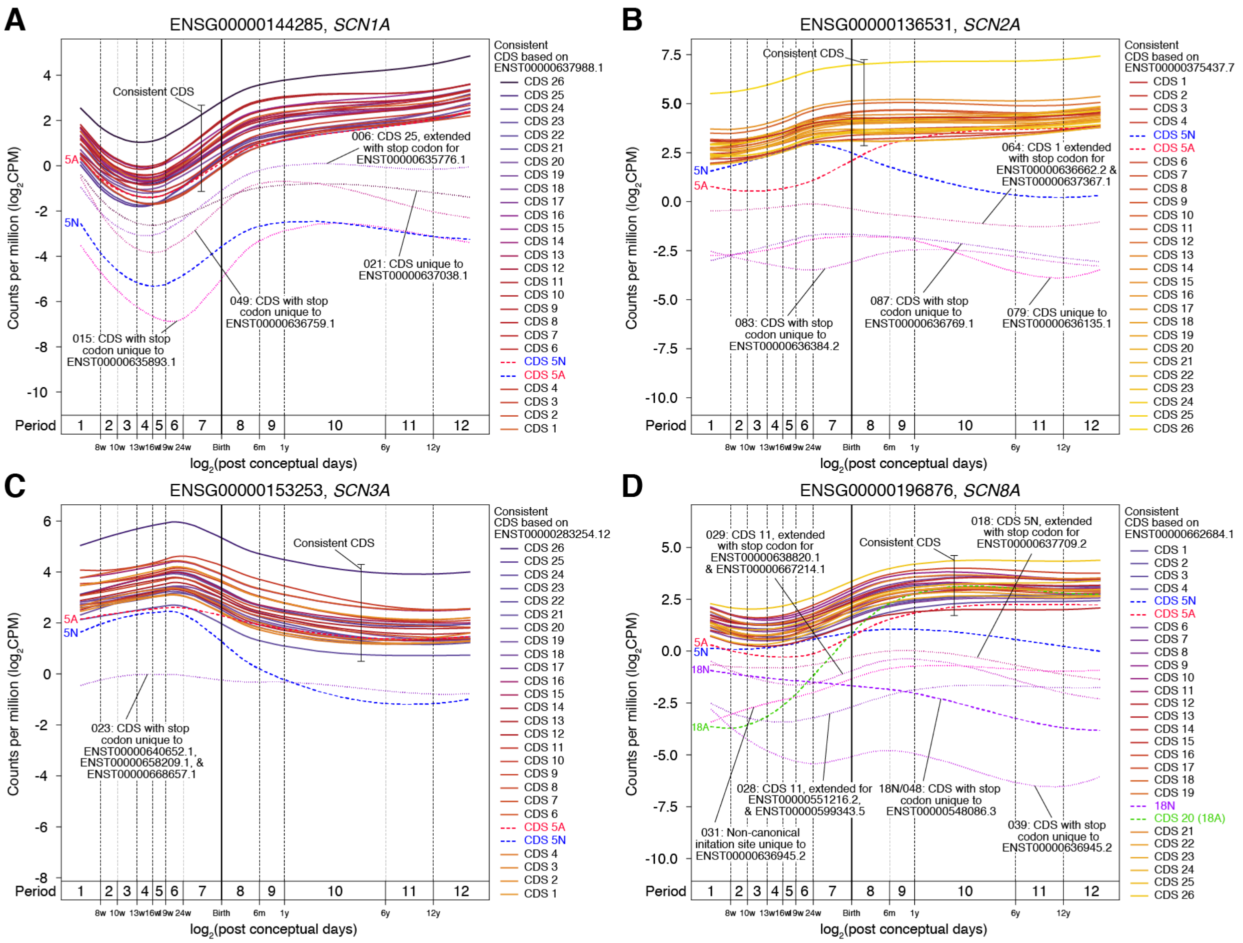


**Figure S3. Expression of all protein-coding exons in voltage-gated sodium channels in the human cortex across development.** **A)** The Loess smoothed average expression of each protein-coding exon in *SCN1A* is shown across development. Constitutive protein-coding exons (CDS) are labelled together and their corresponding color shown in the legend. Variable CDS, that are unique to a few isoform definitions, are labeled along with their DEXSeq exon code (Table S2). **B-D)** Panel 'A' is repeated for *SCN2A*, *SCN3A*, and *SCN8A*.

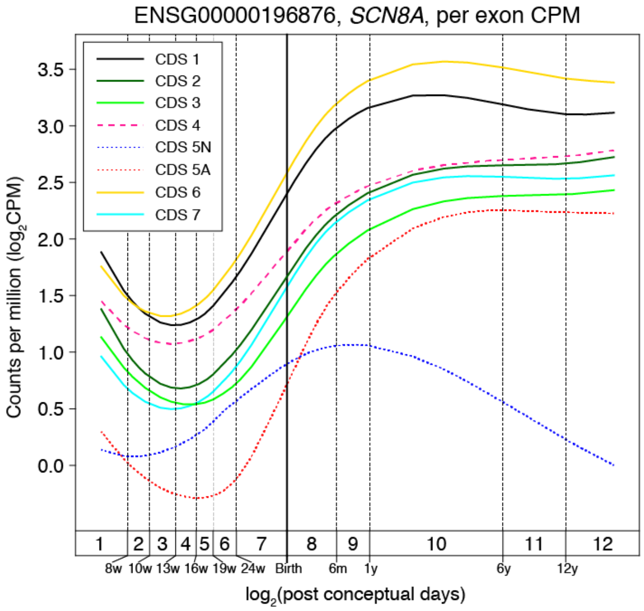


**Figure S4. Expression of *SCN8A* constitutive CDS 1-7 in the human cortex.** The Loess smoothed average expression in the dorsolateral prefrontal cortex is shown across human development. CDS 4 is highlighted in red with a dashed line. These data are a subset of Fig. S2D.

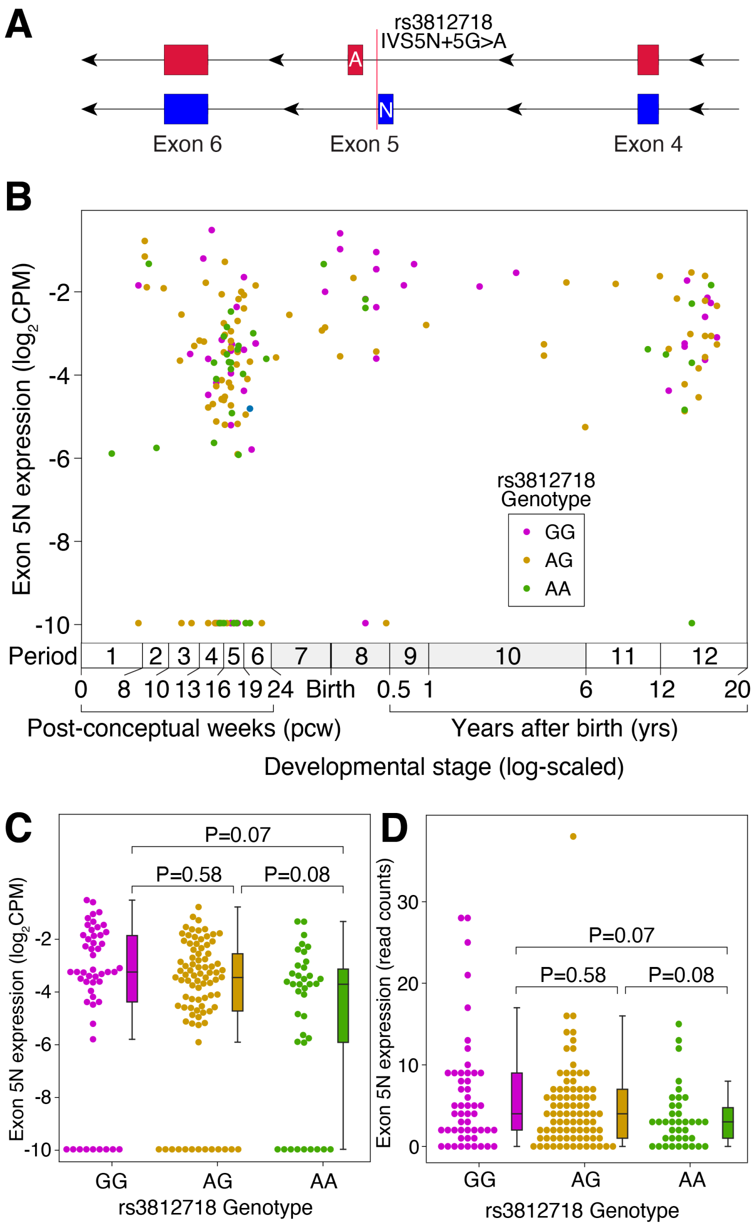


**Figure S5. 5N expression by rs3812718 genotype in the developing human cortex. A)** rs3812718 (chr2:166,053,034 C>T, GRCh38) is a common single nucleotide polymorphism (SNP) previously associated with differential 5N inclusion in the human temporal cortex.[21,22] The SNP is five nucleotides upstream of the 5N donor splice site. **B)** Normalized 5N expression by rs3812718 genotype in 176 *post mortem* human dorsolateral prefrontal cortex samples across cortical development. Expression values are $log_2$-scaled counts per million ($log_2$CPM) with a value of -10 representing no reads detected (31 samples). **C)** Normalized 5N expression ($log_2$CPM) is shown for the same samples as in 'B' binned by genotype. **D)** Raw read counts for 5N expression is shown for the same samples as in 'B' binned by genotype. Statistical tests: C, D) Two-tailed Wilcoxon test.
